# Rapidly predicting vancomycin resistance of *Enterococcus faecium* through MALDI-TOF MS spectrum obtained in real-world clinical microbiology laboratory

**DOI:** 10.1101/2020.03.13.990978

**Authors:** Hsin-Yao Wang, Ko-Pei Lu, Chia-Ru Chung, Yi-Ju Tseng, Tzong-Yi Lee, Jorng-Tzong Horng, Tzu-Hao Chang, Min-Hsien Wu, Ting-Wei Lin, Tsui-Ping Liu, Jang-Jih Lu

**Affiliations:** Department of Laboratory Medicine, Chang Gung Memorial Hospital at Linkou, Taoyuan City, Taiwan; Ph.D. Program in Biomedical Engineering, Chang Gung University, Taoyuan City, Taiwan; Graduate Program in Biomedical Information, Yuan-Ze University, Taoyuan City, Taiwan; Department of Computer Science and Information Engineering, National Central University, Taoyuan City, Taiwan; Department of Information Management, Chang Gung University, Taoyuan City, Taiwan; School of Science and Engineering, The Chinese University of Hong Kong, Shenzhen, China; Warshel Institute for Computational Biology, The Chinese University of Hong Kong, Shenzhen, China; Department of Bioinformatics and Medical Engineering, Asia University, Taichung City, Taiwan; Graduate Institute of Biomedical Informatics, Taipei Medical University, Taipei City, Taiwan; Clinical Big Data Research Center, Taipei Medical University Hospital, Taipei City, Taiwan; Graduate Institute of Biomedical Engineering, Chang Gung University, Taoyuan City, Taiwan; Division of Haematology/Oncology, Department of Internal Medicine, Chang Gung Memorial Hospital at Linkou, Taoyuan City, Taiwan; Biosensor Group, Biomedical Engineering Research Center, Chang Gung University, Taoyuan City, Taiwan; School of Medicine, Chang Gung University, Taoyuan City, Taiwan; Department of Medical Biotechnology and Laboratory Science, Chang Gung University, Taoyuan City, Taiwan

**Author notes:** To whom correspondence should be addressed: JJ Lu **CORRESPONDENCE**: Jang-Jih Lu, M.D., Ph.D., Department of Laboratory Medicine, Chang Gung Memorial Hospital at Linkou, Department of Medical Biotechnology and Laboratory Science, Chang Gung University, 5 Fu-Shing St. Kweishan, Taoyuan 333, Taiwan.

**Keywords:** Vancomycin-resistant *Enterococcus faecium* (VRE*fm*), Antibacterial drug resistance, Matrix-assisted laser desorption ionization time-of-flight (MALDI-TOF) mass spectrometry, Machine learning, Rapid detection

## Abstract

*Enterococcus faecium* is one of the leading pathogens in the world. In this study, we proposed a strategy to rapidly and accurately distinguish vancomycin-resistant *Enterococcus faecium* (VRE*fm*) and vancomycin-susceptible *E. faecium* (VSE*fm*) to help doctors correctly determine the use of vancomycin by a machine learning (ML)-based algorithm. A predictive model was developed and validated to distinguish VRE*fm* and VSE*fm* by analyzing MALDI-TOF MS spectra of unique *E. faecium* isolates from different specimen types. Firstly, 5717 mass spectra, including 2795 VRE*fm* and 2922 VSE*fm*, were used to develop the algorithm. And 2280 mass spectra of isolates, namely 1222 VRE*fm* and 1058 VSE*fm*, were used to externally validate the algorithm. The random forest-based algorithm demonstrated good classification performances for overall specimens, whose mean AUROC in 5-fold cross validation, time-wise validation, and external validation was all greater than 0.84. For the detection of VRE*fm* in blood, sterile body fluid, urinary tract, and wound, the AUROC in external validation was also greater than 0.84. The predictions with algorithms were significantly more accurate than empirical antibiotic use. The accuracy of antibiotics administration could be improved by 30%. And the algorithm could provide rapid antibiotic susceptibility results at least 24 hours ahead of routine laboratory tests. The turn-around-time of antibiotic susceptibility could be reduced by 50%. In conclusion, a ML algorithm using MALDI-TOF MS spectra obtained in routine workflow accurately differentiated VRE*fm* from VSE*fm*, especially in blood and sterile body fluid, which can be applied to facilitate the clinical testing process due to its accuracy, generalizability, and rapidness.

## Introduction

*Enterococcus* spp. is one of the leading pathogens in healthcare-associated infection.^1^ Enterococcal infection could cause urinary tract infection, blood stream infection, and even mortality.^2^ Until recently, vancomycin was virtually the only drug that could be consistently relied on for treating multidrug-resistant enterococcal infections^3,4^. Vancomycin-resistant *Enterococcus* (VRE) has led to heavy burden on healthcare worldwide since its first-time isolation.^5,6^ *Enterococcus faecalis* and *E. faecium* are the 2 most commonly isolated *Enterococcus* spp. in clinical practice.^1^ VRE *faecium* (VRE*fm*) has received considerably more attention than VRE *faecalis* (VRE*fs*) because most of the clinically isolated VRE is *E. faecium* in the recent decades^4,7^ and VRE*fm* causes more severe infection than VRE*fs*^8,9^. Early detection of vancomycin resistance is essential for successfully treating VRE*fm* infection.^10^ Vancomycin could be discontinued, and antimicrobial agents could be replaced with other antibiotics (eg, linezolid and daptomycin) based on the laboratory results of vancomycin resistance^11,12^. Patients’ prognosis could be improved and further drug resistance development could be avoided by using susceptible antibiotics.^11^ However, typical tests in clinical microbiology laboratories, such as the minimal inhibitory concentration test or agar-diffusion test, fail to provide results for antibiotic susceptibility rapidly. The antibiotic susceptibility test (AST) of vancomycin is time-consuming, and the Clinical and Laboratory Standards Institute recommended a full 24 hours should be held for accurate detection of vancomycin resistance in enterococci.^13^ This would considerably delay accurate prescription of antibiotics against *E. faecium*. Furthermore, prescribing antibiotics based on empirical prescription, without determining AST, would result in low effectiveness (approximately 50%), depending on the local epidemiology of VRE*fm*.^12^ Thus, a new tool is needed to provide AST for VRE*fm* rapidly and accurately.

Recently, matrix-assisted laser desorption ionization time-of-flight (MALDI-TOF) mass spectrometry (MS) has become popular among clinical microbiology laboratories worldwide because of its reliability, rapidity, and cost-effectiveness in identifying bacterial species.^14–16^ In addition to species identification, MALDI-TOF MS has been promising in other applications, such as strain typing or AST.^17–19^ MALDI-TOF MS can generate massive data comprising hundreds of peak signals on the spectra.^17,20^ The complex data of MALDI-TOF spectra are overwhelming to even an experienced medical staff.^19^ Studies have attempted to identify the characteristic peak through visual inspection.^21,22^ The results of the studies have been discordant, which has limited the clinical utility.^23–25^

Machine learning (ML) is a good analytical method for solving classification problems through identification of implicit data patterns from complex data.^26^ The ML method outperforms traditional statistical methods because of its excellent ability to handle complex interactions between large amount of predictors and good performance in non-linear classification problems^27^ ML has been successfully applied in several clinical fields.^27–36^ Thus, the ML algorithm is especially appropriate for analyzing complex data such as MALDI-TOF spectra. However, to our knowledge, few studies have used ML in the analysis of MALDI-TOF spectra for rapidly reporting VRE*fm*, and the case numbers in these studies were insufficient, and so, ML algorithm generalization has been limited.^37–39^ Moreover, to date, no study has validated AST prediction ML algorithms by using large real-world data.

In this study, we aimed to develop and validate a VRE*fm* prediction ML model by using consecutively collected real-world data from 2 tertiary medical centers (Chang Gung Memorial Hospital [CGMH], Linkou branch and CGMH, Kaohsiung branch). Using the largest MALDI-TOF spectrum clinical data to date, the ML algorithm could predict VRE*fm* accurately, rapidly, and in a ready-to-use manner based on the real-world evidence, which is more representative for clinical practice.^40^ Moreover, we confirmed the robustness and generalization of the ML algorithm through several validation methods, namely cross-validation, time-wise internal validations (unseen independent testing dataset classified according to time), and external validation (unseen independent testing dataset from another medical center). According to the real-world evidence-based validation, our VRE*fm* prediction ML models are ready to be incorporated into routine workflow.

## Materials and methods

### Data source

We designed a novel machine learning approach which can improve accuracy of antibiotics administration and reduce the turn-around-time of antibiotics susceptibility test. We summarized the comparison between the machine learning approach and the traditional approach used in current clinical microbiology laboratory. We schematically illustrated the study design in Figure 1(b). The data used in this retrospective study was consecutively collected from the clinical microbiology laboratories of 2 tertiary medical centers in Taiwan, namely CGMH Linkou branch and CGMH Kaohsiung branch between January 1, 2013 and December 31, 2017. The clinical microbiology laboratories collected and processed all the routine specimens obtained from the hospitals. In total, 7997 *E. faecium* cases were identified and included in this study, whereas 5717 (VRE*fm*: 48.89%) and 2280 (VRE*fm*: 53.60%) cases, respectively, were obtained from Linkou and Kaohsiung branches of CGMH. The *E. faecium* strains were isolated from blood, urinary tract, sterile body fluids, and wound. The detailed description of specimen types is provided in eTable 1 in the Supplement. The study was approved by the Institutional Review Board of Chang Gung Medical Foundation (No. 201900767B0). We followed the Standards for Reporting of Diagnostic Accuracy 2015^41^ and the Transparent Reporting of a Multivariable Prediction Model for Individual Prognosis or Diagnosis reporting guidelines.^42^

**Figure 1(a).**
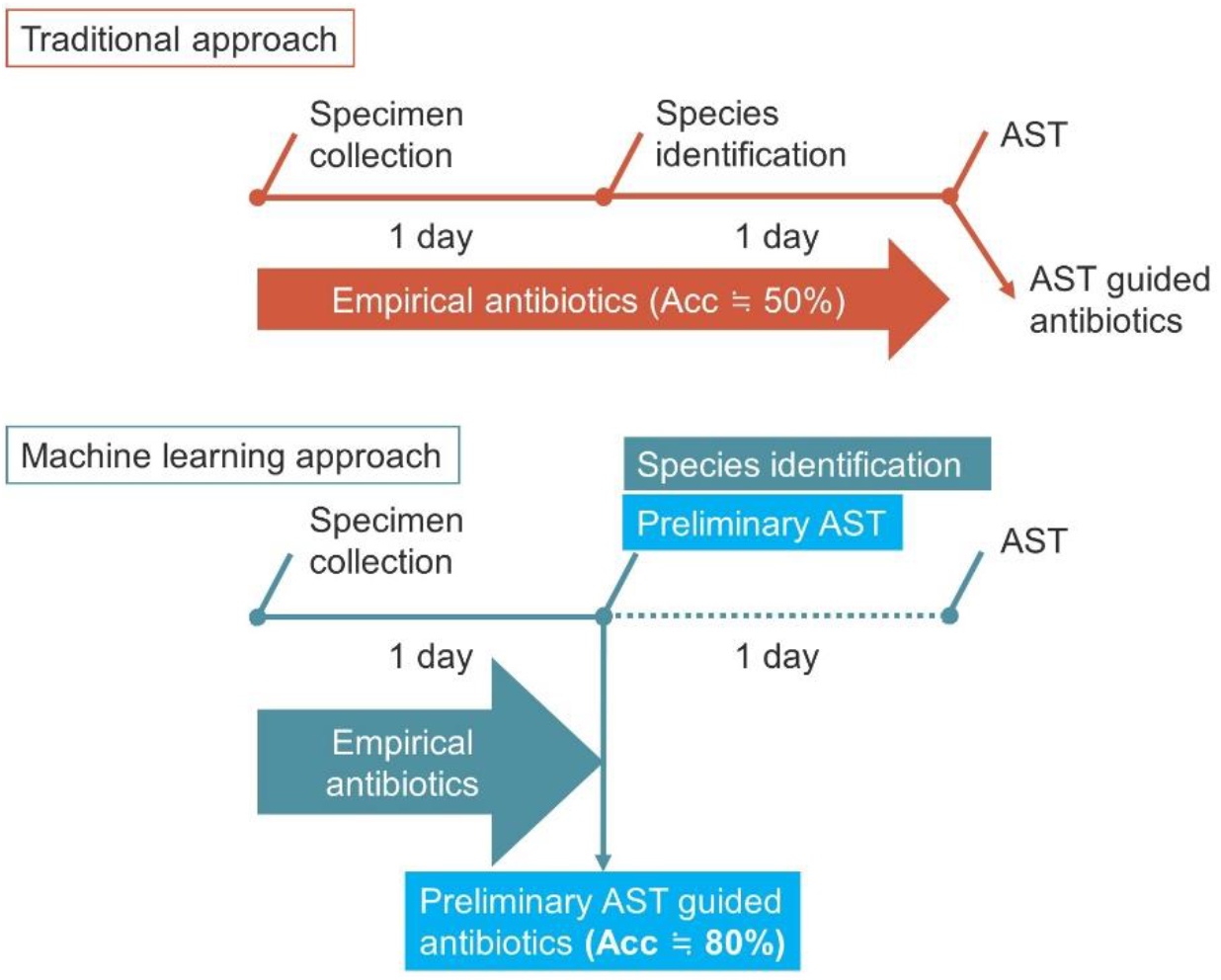
Scheme of using the VRE*fm* Model. We plotted a timeline of bacterial culture test in current clinical microbiology laboratory (i.e., traditional approach) and a modified timeline when the VRE*fm* model is incorporated (i.e., machine learning approach). In the traditional approach, specimens are collected for bacterial culture test. One day is usually needed for growth of a single colony for species identification (by MALDI-TOF MS). Antibiotics susceptibility test (AST) of vancomycin for VRE*fm* will cost another day to report. By contrast, in the machine learning approach, the VRE*fm* model can provide preliminary AST at the time when bacterial species is identified by MALDI-TOF MS. For treating VRE*fm*, the machine learning approach can improve accuracy of antibiotics use by around 30% (from 50% accuracy of empirical antibiotics use in the traditional approach to 80% accuracy of preliminary AST provided by the machine learning approach). Meanwhile, the turn-around-time of bacterial culture test can be reduced to one day, which is 50% reduction.

**Figure 1(b).**
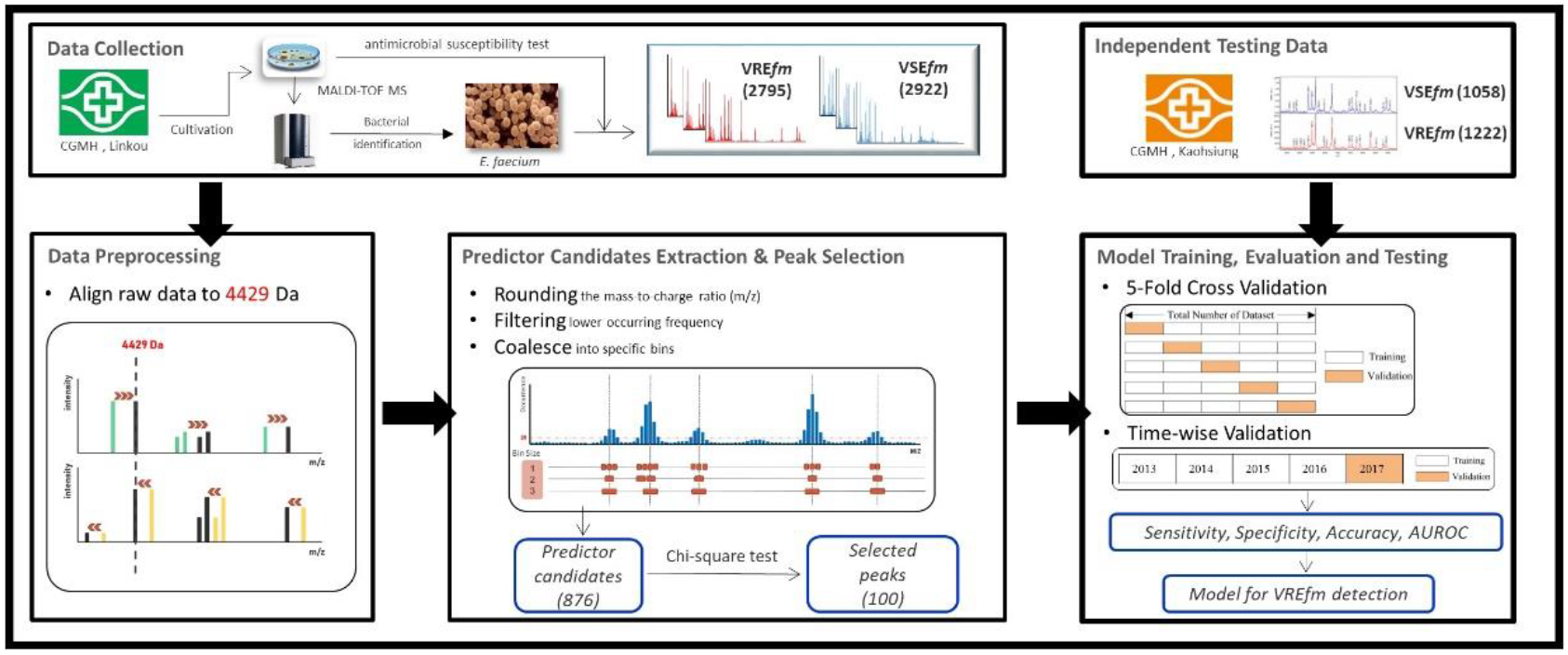
Schematic Illustration of the Study Design. We developed and validated a VRE*fm* prediction model. The study included several steps, namely data collection, data preprocessing, predictor candidate extraction and important predictor selection, model training, evaluation, and testing. In data collection, data were obtained from 2 tertiary medical centers (Linkou and Kaohsiung branches of CGMH). The data included mass spectra and results of the vancomycin susceptibility test of *E. faecium*. Data from the CGMH Linkou branch were used for model training and validation, while data from the CGMH Kaohsiung branch served as an independent testing data. In the steps of data preprocessing and predictor candidate extraction and important predictor selection, a specific set of crucial predictors would be used for model training. K-fold, time-wise CV, and external validation were used to confirm the models’ robustness. The VRE*fm* prediction model can detect VRE*fm* accurately at least 1 day earlier than the current method.

### Definition of *E. faecium* and vancomycin susceptibility

*E. faecium* was identified using MALDI-TOF spectra measured using a Microflex LT mass spectrometer and analyzed using Biotyper 3.1 (Bruker Daltonik GmbH, Bremen, Germany). A log score (generated through Biotyper 3.1) larger than 2 was used for confirming *E. faecium*.^17–19^ We tested vancomycin susceptibility of *E. faecium* by using the paper disc method. The details of *E. faecium* identification and AST are given in the eMethods in the Supplement.

### MALDI-TOF mass spectrum data collection and preprocessing

The details were described in the Supplements.

### Peak selection from MALDI-TOF mass spectra for model development

We applied the embedded feature-selection method to select the most important peaks from MALDI-TOF mass spectra.^43^ The peaks were ranked using the p-values of the chi-square test of homogeneity, which was employed to determine whether frequency counts were distributed identically across VRE*fm* and vancomycin-susceptible *E. faecium* (VSE*fm*). Preliminarily, we selected top 10 important peaks to plot a heat map based on the hierarchical clustering (eMethods in the Supplement). All the ranked peaks were incorporated in the model accordingly until the performance did not increase. Consequently, we could obtain the important peaks that were highly related to differentiation of VRE*fm* and VSE*fm* isolates.

For determining the number of peaks included in the ML models, we forwardly added them into the ML models and calculated the performance using accuracy as the metric. First, the predictor candidates were sorted in a descending order according to the importance score, and one predictive peak was added at a time into the ML models. On the basis of predictive peak composition, we used different algorithms, namely random forest (RF), support vector machine (SVM) with a radial basis function kernel, and k-nearest neighbor (KNN) and applied 5-fold cross validation (CV) in the data from the CGMH Linkou branch. The accuracies of the ML models were calculated to determine the adequate number of predictive peaks included in the models.

### Development and validation of VRE*fm* prediction models

We aimed to develop and validate a robust VRE*fm* prediction model capable of detecting VRE*fm* earlier than the AST report. Three commonly used ML algorithms, namely RF, SVM with a radial basis function kernel, and KNN, were used for developing the VRE*fm* prediction model. These ML algorithms have demonstrated their successful applications (either classification or prediction) in clinical practice.^17–19,27,28,35,36^ The details of these ML algorithms and model training processes are attached in the eMethods in the Supplement.

We thoroughly evaluated the performance and robustness of the VRE*fm* prediction models using 5-fold CV, time-wise internal validation, and external validation. Data from the CGMH Linkou branch were used for 5-fold CV and time-wise internal validation; by contrast, data from the CGMH Kaohsiung branch served as the unseen independent testing data for external validation. For 5-fold CV, data were randomly divided into 5 datasets. Each one of the 5 datasets served as the testing dataset to evaluate the performance of the model developed by the other 4 datasets. In 5-fold CV, we obtained 5 measurements of metrics for evaluating the robustness of VRE*fm* prediction models. Moreover, to evaluate performance using prospectively collected data, we conducted time-wise internal validation: we used data collected between January 1, 2013 and December 31, 2016 as the training dataset for developing VRE*fm* prediction models, while data from January 1, 2017 to December 31, 2017 served as the testing dataset. To test the generalizability of the models, we used data from the CGMH Linkou branch to develop VRE*fm* prediction models and used data from the CGMH Kaohsiung branch to test the models’ performance in a different institute. Additionally, we evaluated the performance of the VRE*fm* prediction model using different types of specimens, namely blood, urinary tract, sterile body fluid, and wound, by using data from the CGMH Kaohsiung branch. We adopted metrics including sensitivity, specificity, accuracy, positive predictive value (PPV), negative predictive value (NPV), receiver operating characteristic (ROC) curve, and area under the receiver operating characteristic curve (AUROC) to access and compare the performance of the VRE*fm* prediction model.

### Statistical analysis

The confidence intervals for sensitivity, specificity, and accuracy were estimated using the calculation of the confidence interval for a proportion in one sample situation. Specifically, the critical values followed the Z-score table. To compare the percentages in matched samples, Cochran’s Q test, a nonparametric approach, was implemented in this study.^44^ Then, we employed pairwise McNemar’s tests^45^ for post hoc analysis and adopted the false discovery rate proposed by Benjamini and Hochberg (1995) to adjust the *P* value.^46^ Furthermore, the confidence intervals of AUROCs were determined using the nonparametric approach, and the AUROC comparisons mainly adopted the nonparametric approach proposed by Delong et al.^47^

## Results

### Predictive peaks for detecting VRE*fm*

We defined crucial predictive peaks when the occurrence frequency of a peak was significantly different (defined by the chi-square test) in VRE*fm* and VSE*fm*. In the step of extracting predictor candidates, 876 predictor candidates were extracted. From the predictor candidates, we used the chi-square method to select important predictive peaks.

We selected 10 most critical predictive peaks and plotted a heat map to preliminarily visualize the difference between VRE*fm* and VSE*fm* (Figure 2). Peaks of *m/z* 3172, 3302, 3645, 6342, 6356, 6603, and 6690 were found more frequently in VRE*fm*; by contrast, *m/z* 3165, 3681, and 7360 occurred more frequently in VSE*fm*. Although these important predictive peaks were statistically significant, we found them in both VRE*fm* and VSE*fm*. The full list of crucial predictive peaks is provided in eTable 2 in the Supplement.

**Figure 2.**
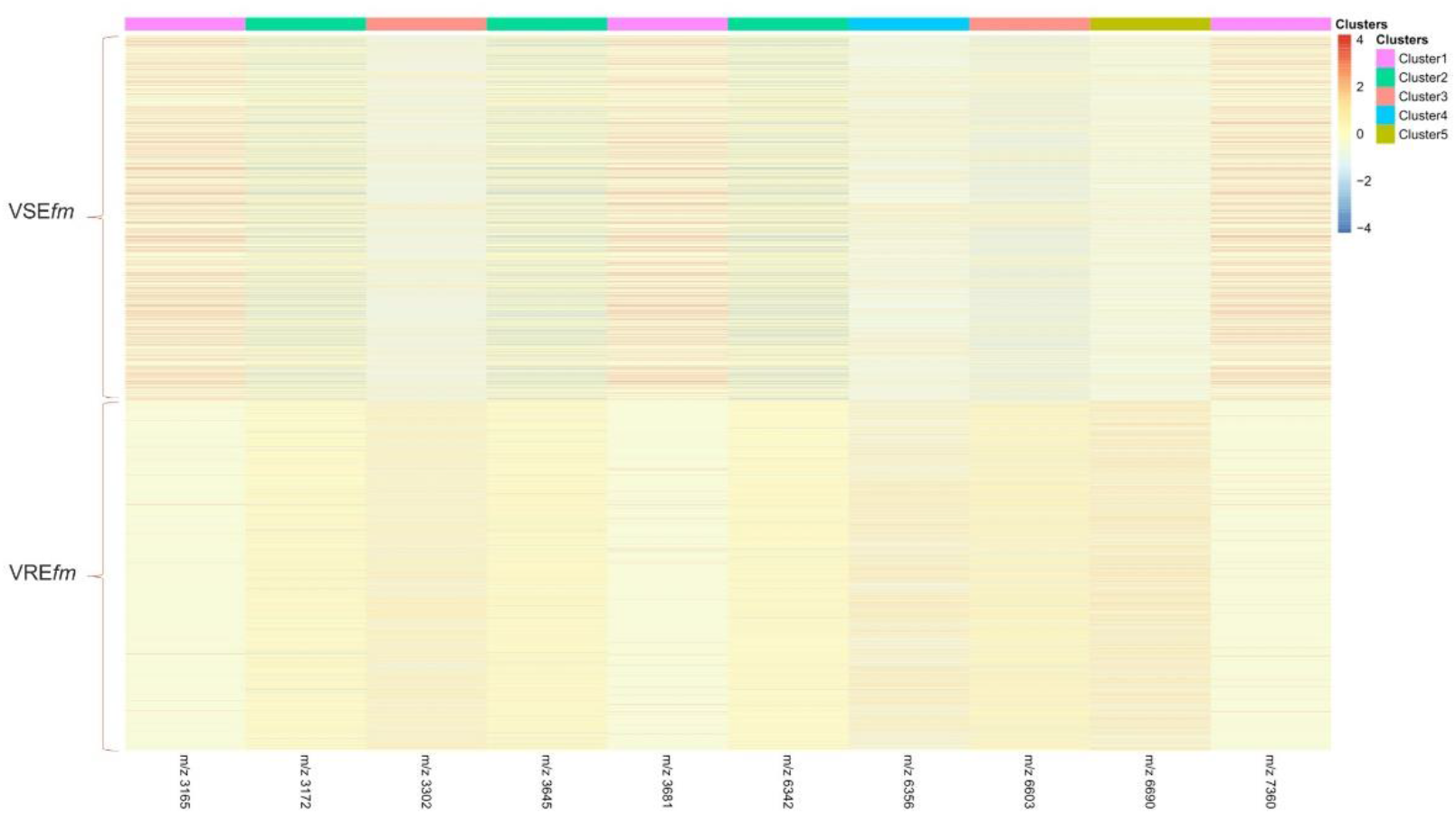
Heat map. We selected top 10 discriminative peaks by chi-square testing the occurrence frequency of peaks in VRE*fm* and VSE*fm*. The heat map was plotted based on the hierarchical clustering of all the VRE*fm* and VSE*fm* isolates from the CGMH Linkou branch. Rows represent the isolates, and columns represent the top 10 discriminative peaks. The values in the heat map represent the MS spectral intensity which was log_10_-normalized and z-score standardized. Red color indicates relatively higher peak intensity while blue color indicates lower peak intensity. The isolates are grouped into 5 clusters. VRE*fm* and VSE*fm* isolates can be visually differentiated by using the top 10 discriminative peaks.

We selected several important predictive peaks from the predictor candidate list, which was ordered according to the chi-square score. eFigure 4 in the Supplement shows the change in ML models performance when the number of critical predictive peaks increased. For all the ML algorithms used in the study, a similar trend of performance was observed: the accuracies of the ML models reached a steady plateau when the included number of important predictive peaks was larger than 100 (eFigure 4 in the Supplement). Thus, the top 100 crucial predictive peaks were selected as the peak composition for the following experiments.

### Performance of VRE*fm* prediction models

We summarized the ML models’ performance in Table 1, Table 2, and Figure 3. The details of comparison between different algorithms are described in the Supplement. The RF model outperformed SVM and KNN in 5-fold CV, time-wise internal validation, and external validation (eTable 3 in the Supplement), where the AUROC ranged from 0.8463 to 0.8553 and accuracy ranged from 0.7769 to 0.7855. Moreover, performance robustness was also observed in SVM and KNN. Figure 3 shows typical ROC curves for the 3 algorithms in all the 3 validations. We used Youden’s index to select the threshold from the ROC curves in search of balanced sensitivity and specificity. In external validation, the sensitivity and specificity of RF were 0.7791 (95% confidence interval: 0.7620-0.7961) and 0.7930 (95% confidence interval: 0.7764-0.8096). On the basis of the resistance rate (VRE*fm*: 53.60%) in the external validation dataset, the PPV was 0.8130 and the NPV was 0.7565.

**Table 1.** Performance of VRE*fm* Prediction Models in Terms of k-Fold CV, Time-Wise Validation, and External Validation.

| AUROC | RF | SVM | KNN |
| --- | --- | --- | --- |
| 5-fold CV | 0.8495 (0.8397, 0.8594) | 0.8367 (0.8264, 0.8471) | 0.7908 (0.7792, 0.8024) |
| Time-wise validation | 0.8463 (0.8273, 0.8654) | 0.8368 (0.8169, 0.8566) | 0.7908 (0.7690, 0.8127) |
| External validation | 0.8553 (0.8399, 0.8706) | 0.8407 (0.8246, 0.8569) | 0.8050 (0.7872, 0.8227) |
| Accuracy |  |  |  |
| 5-fold CV | 0.7769 (0.7660, 0.7878) | 0.7610 (0.7499, 0.7721) | 0.7248 (0.7131, 0.7364) |
| Time-wise validation | 0.7840 (0.7640, 0.8039) | 0.7815 (0.7615, 0.8016) | 0.7228 (0.7011, 0.7445) |
| External validation | 0.7855 (0.7687, 0.8024) | 0.7781 (0.7610, 0.7951) | 0.7355 (0.7174, 0.7536) |
| Sensitivity |  |  |  |
| 5-fold CV | 0.8054 (0.7951, 0.8517) | 0.7826 (0.7719, 0.7934) | 0.7873 (0.7767, 0.7980) |
| Time-wise validation | 0.8153 (0.7965, 0.8341) | 0.8415 (0.8238, 0.8592) | 0.7491 (0.7281, 0.7702) |
| External validation | 0.7791 (0.7620, 0.7961) | 0.7954 (0.7789, 0.8120) | 0.8044 (0.7881, 0.8207) |
| Specificity |  |  |  |
| 5-fold CV | 0.7497 (0.7384, 0.7609) | 0.7403 (0.7289, 0.7517) | 0.6649 (0.6526, 0.6772) |
| Time-wise validation | 0.7477 (0.7266, 0.7688) | 0.7120 (0.6900, 0.7340) | 0.6922 (0.6698, 0.7146) |
| External validation | 0.7930 (0.7764, 0.8096) | 0.7580 (0.7405, 0.7756) | 0.6560 (0.6365, 0.6755) |

AUROC, area under the receiver operating characteristic curve.

**Table 2.**
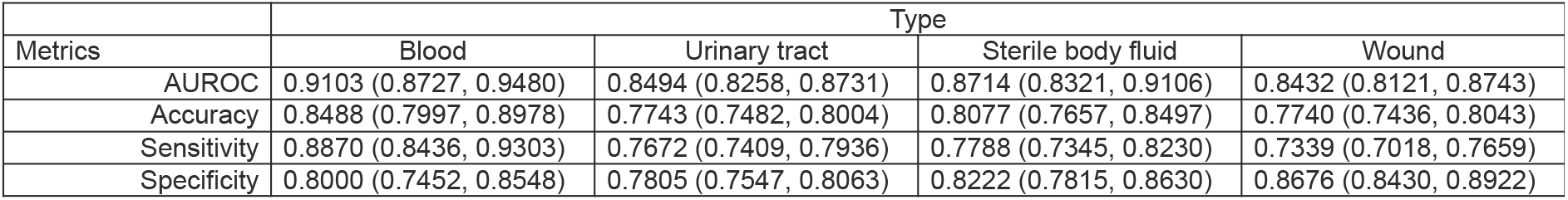
Performance of the RF-Based VRE*fm* Detection Model With Different Types of Specimens in Terms of External Validation.

| Metrics | Type |  |  |  |
| --- | --- | --- | --- | --- |
|  | Blood | Urinary tract | Sterile body fluid | Wound |
| AUROC | 0.9103 (0.8727, 0.9480) | 0.8494 (0.8258, 0.8731) | 0.8714 (0.8321, 0.9106) | 0.8432 (0.8121, 0.8743) |
| Accuracy | 0.8488 (0.7997, 0.8978) | 0.7743 (0.7482, 0.8004) | 0.8077 (0.7657, 0.8497) | 0.7740 (0.7436, 0.8043) |
| Sensitivity | 0.8870 (0.8436, 0.9303) | 0.7672 (0.7409, 0.7936) | 0.7788 (0.7345, 0.8230) | 0.7339 (0.7018, 0.7659) |
| Specificity | 0.8000 (0.7452, 0.8548) | 0.7805 (0.7547, 0.8063) | 0.8222 (0.7815, 0.8630) | 0.8676 (0.8430, 0.8922) |

**Figure 3(a).**
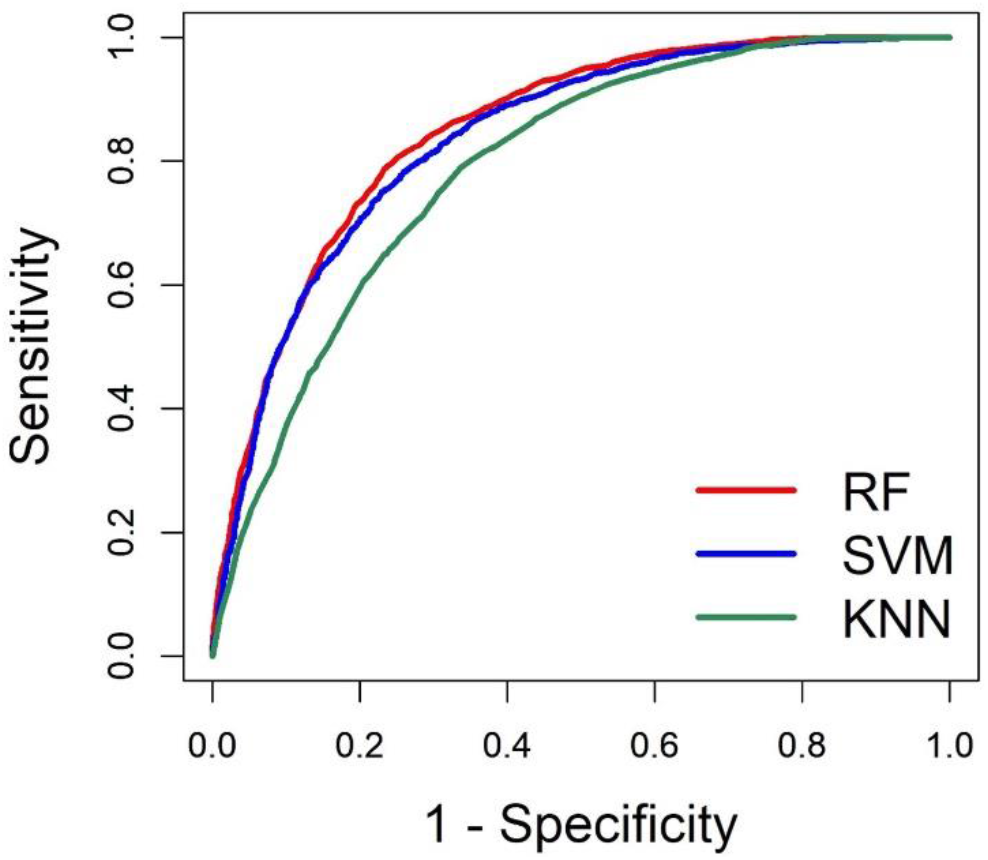
ROC Curves for Different Algorithms in Terms of Linkou 5-Fold CV.

**Figure 3(b).**
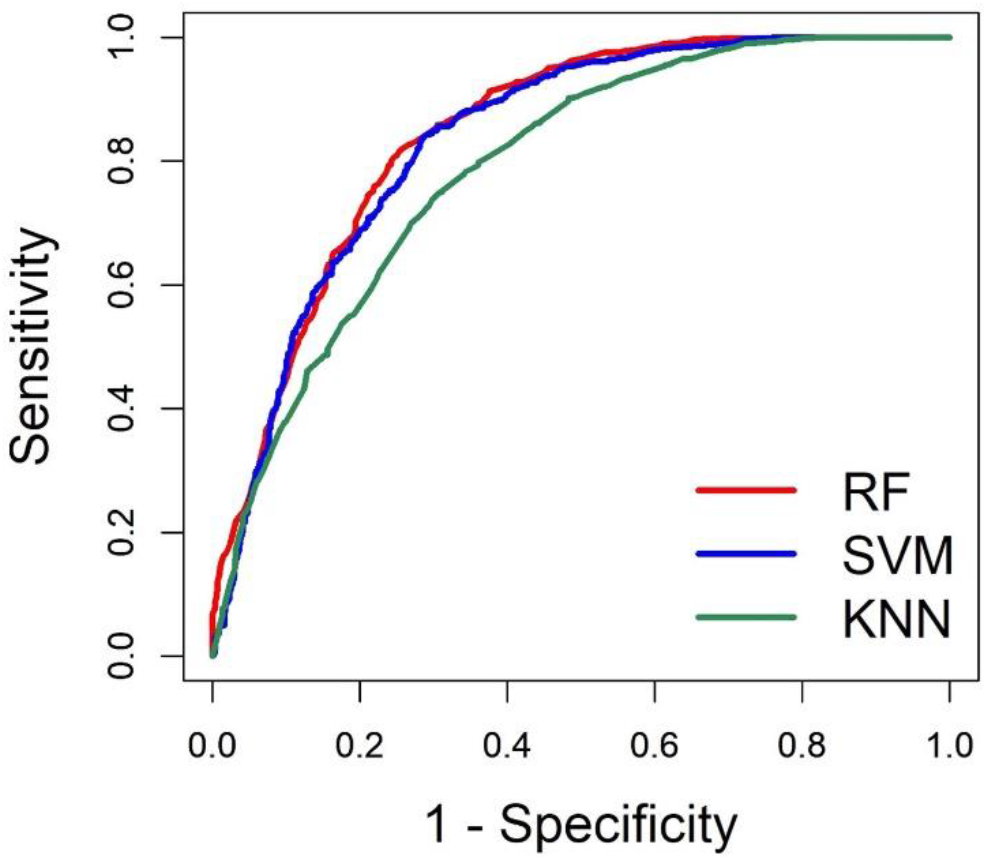
ROC Curves for Different Algorithms in Terms of Time-Wise Validation.

**Figure 3(c).**
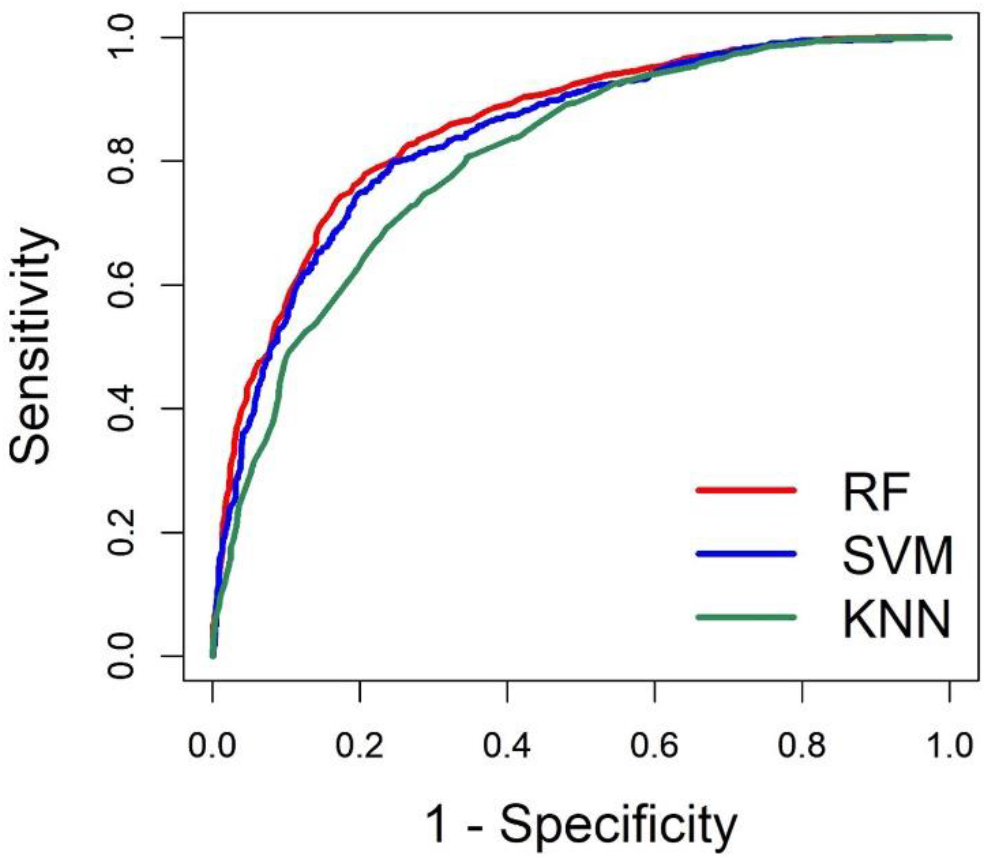
ROC Curves for Different Algorithms in Terms of External Validation.

**Figure 3(d).**
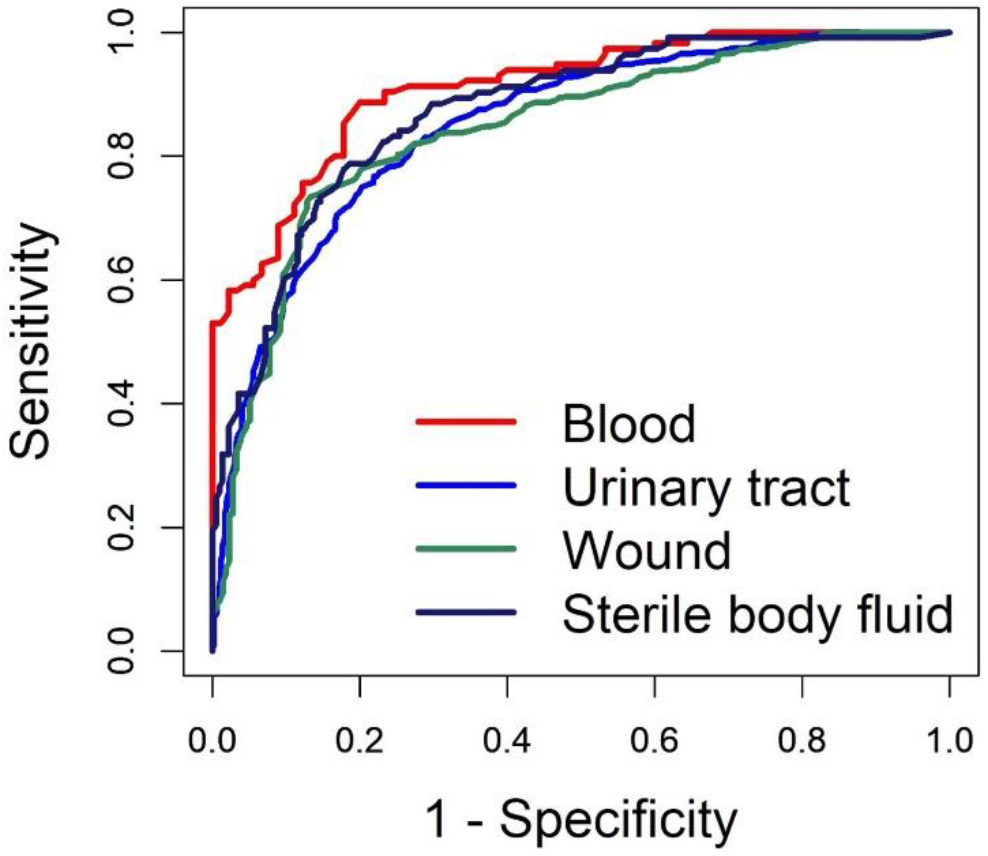
ROC Curves for the RF-Based VRE*fm* Model With Different Types of Specimens.

Given that the RF algorithm attained the highest performance, additionally, we tested the performance of the RF-based VRE*fm* prediction model using different types of specimens in the independent testing dataset (ie, external validation by using data of the CGMH Kaohsiung branch) (Table 2). The RF-based VRE*fm* prediction model attained higher performance in predicting VRE*fm* in blood and sterile body fluid specimens than the other specimen types. The AUROC of blood specimens reached 0.9103 (95% confidence interval: 0.8727-0.9480), whereas that of sterile body fluid specimens reached 0.8714 (95% confidence interval: 0.8321-0.9106). Moreover, the sensitivity (0.8870, 95% confidence interval: 0.8436-0.9303) and specificity (0.8000, 95% confidence interval: 0.7452-0.8548) of the RF-based VRE*fm* prediction model for the blood specimen were also balanced and significantly higher than those for other specimens. By contrast, the performance of the RF-based VRE*fm* prediction model for urinary tract specimens (0.8494, 95% confidence interval: 0.8258-0.8731) was similar to that for overall specimens (0.8553, 95% confidence interval: 0.8399-0.8706).

## Discussion

We developed ML-based models for predicting VRE*fm* rapidly and accurately based on MALDI-TOF MS data. The models were especially effective in predicting VRE*fm* in invasive infections (ie, blood and sterile body fluid). We used the largest up-to-date real-world data to validate the robustness and generalization of the ML-based models by using k-fold CV, time-wise internal validation, and external validation. The rapid and accurate AST of vancomycin is promising for determining antibiotics against VRE*fm* infection.

Our results suggested that AST could be predicted accurately by using ML algorithms to analyze MALDI-TOF MS data. MALDI-TOF MS is a powerful analytical tool in current clinical microbiology laboratories because of its rapidness and cost-effectiveness in identifying bacterial species.^14–16^ On the basis of the massive data produced by MALDI-TOF MS, moreover, some studies have demonstrated that subspecies typing could be predicted from a specific pattern of MS spectra only.^17,19^ Furthermore, other studies have shown a good correlation between AST and specific patterns of MS spectra.^18,23–25,48^ However, some issues have limited the generalization of these results. First, most of the studies have adopted an additional protein extraction step before analytical measurement of MALDI-TOF MS. The protein extraction step could enhance data quality; however, it is not routinely used in clinical practice because it is labor-intensive, time-consuming, and expensive.^17,18^ By contrast, we used the direct deposition method, which is recommended by the manufacturer and is used for everyday works. Thus, our models are more feasible for the existing workflow because they were trained using real-world data. Second, the data sizes in these studies were too small to be representative. We demonstrated that the ML-based models for predicting VRE*fm* can be applied as a clinical decision support tool by using the largest up-to-date datasets collected through the direct deposition method and various validation methods.

Identifying crucial predictive peaks in VRE*fm* classification may not be essential in clinical application; however, the specific combination of crucial predictive peaks would inspire further studies investigating the molecular mechanism of VRE*fm*. Typically, the *vanA* cluster is the most common mediator of vancomycin resistance in enterococci,^49^ although many vancomycin resistance genes have been identified.^50^ In brief, many factors together attribute to antibiotic resistance. Moreover, the complex mechanisms of antibiotic resistance would evolve in response to the selective pressures of their competitive environment (eg, antibiotic use).^49^ Thus, identifying the important predictive peaks for VRE*fm* could help us understand the mechanism behind resistance. In this study, for example, peaks of *m/z* 6603, 6631, and 6635 were found frequently for VRE*fm* (eTable 2 in the Supplement). The finding is consistent with a previous study where Griffin et al. reported *m/z* 6603 is specific for *vanB*-positive VRE*fm*, while *m/z* 6631 and 6635 are specifically found for *vanA*-positive VRE*fm*.^38^ These peaks are worthy of further identification in future investigations. Moreover, new antibiotics against VRE*fm* can be developed based on these predictive peaks for VRE*fm*.

Our ML models persistently performed well in 5-fold CV, time-wise internal validation, and external validation. Moreover, all the ML algorithms used in this study exhibited good performance (AUROC > 0.8). It could be explained that discriminating VRE*fm* from VSE*fm* is generally achievable after adequate feature extraction and feature selection processes. In time-wise internal validation, we intended to simulate a prospective study for a model trained by the “past data” to analyze the “future data.” Based on the performance of time-wise internal validation, we concluded that the trained ML models could also perform well on the prospectively collected data, which are unseen in the training process. Previous study results differentiating VRE*fm* from VSE*fm* by using MALDI-TOF MS spectra could not be generalized.^23–25,38^ The inconsistent results could be because less features (<10) were used. A review article reported that peak-level reproducibility of MALDI-TOF mass is approximately 80%.^51^ The classification performance is compromised when essential peaks are few and happen to be absent on the mass spectra. In our study, the ML models performed stably when the included peaks were more than 100 (eFigure 4 in the Supplement). The steady and good performance of our ML models could be explained by the much more included peaks: when some of the essential peaks are not reproduced in the mass spectra, we can still use other alternative essential peaks to conduct an accurate classification. The number of essential peaks somehow compensated the insufficient reproducibility of MALDI-TOF mass. By contrast, regarding predicting VRE*fm* for various specimens, we found that the RF-based model performed especially well in blood and sterile body fluids. The superior prediction performance could be attributed to the relatively fewer number of VRE*fm* strains in blood and sterile body fluids. Bacterial infection in blood or sterile body fluids is typically regarded as invasive infection.^52^ Only a few VRE*fm* strains (sequence type (ST)17, ST18, ST78, and ST203) cause invasive infections in blood or sterile body fluids according to the studies in Taiwan^53^ and Ireland.^54^ The nature of the classification problem would be more simple when the number of labels is fewer.

## Limitations

This study has several limitations. First, although the models were evaluated using unseen external data from different medical centers, all the training data and testing data were collected from only 2 tertiary medical centers in Taiwan. Directly applying the ML models in hospitals of other areas or countries as well as in primary care institutes may not be appropriate. However, we believe that the method, but not the trained model, could be generalized. Although our ML models were validated comprehensively using 3 different approaches and the results show that the difference in MALDI-TOF mass spectra between VRE*fm* and VSE*fm* can be distinguished through all the ML algorithms we used, we suggest others collecting their locally relevant data for training and validating the VRE*fm* predicting model given that the epidemiology of VRE*fm* could be fairly different site by site. Second, our primary goal was to develop and validate a practical and ready-to-use ML model in real-world practice. We found some crucial predictive peaks for VRE*fm*; however, we did not confirm the identities for these peaks. It is worthy of identifying these peaks in further investigations. Third, we did not use the deep learning (DL) algorithm for predicting VRE*fm*, although DL has been successful in the image classification or radiology field.^32,33^ In this study, VRE*fm* could be accurately predicted using several classic algorithms (ie, RF, SVM, and KNN) that require less resource and time in training and using models. Moreover, DL usually requires more training samples and is financially and computationally more expensive than classical ML algorithms.^55^ DL utility in analyzing MS data rather than image data could be another promising issue in the bioinformatics field. Fourth, no strain typing data were included. Thus, the molecular epidemiology of VRE*fm* used in this study is unknown.

## Conclusions

We developed and validated robust ML models capable of discriminating VRE*fm* from VSE*fm* based on MALDI-TOF MS spectra. These models were especially good at detecting VRE*fm* causing invasive diseases. The accurate and rapid detection of VRE*fm* by using the ML models would facilitate more appropriate antibiotic prescription.

## Acknowledgments

This manuscript was edited by Wallace Academic Editing.

## Author Contributions

HYW, KPL, and CRC had full access to all the data in the study and take responsibility for the integrity of the data and the accuracy of data analysis. HYW, KPL, CRC, and YJT analyzed/interpreted the data, performed experiments, designed the study, and wrote the manuscript. HYW, CRC, YJT, JTH, TYL, THC, MHW, TPL, and JJL reviewed/edited the manuscript for important intellectual content and provided administrative, technical, or material support. JJL obtained funding and supervised the study.

