## Supplemental materials for "Rapidly predicting vancomycin resistance of *Enterococcus faecium* through MALDI-TOF MS spectrum obtained in real-world clinical microbiology laboratory"

### Specimen processing, *Enterococcus faecium* identification, and vancomycin susceptibility test

Clinical specimens were continuously collected as daily routine from all the wards to the clinical microbiology laboratory of Chang Gung Memorial Hospital, both Linkou and Kaohsiung branches. The specimen types included blood, respiratory tract specimen (*ie,* sputum, bronchial wash, and bronchoalveolar lavage), sterile cavity fluid (*ie*, ascites, pleural effusion, pericardial effusion, cerebrospinal fluid, and synovial fluid), tip of implant, urine, wound, and others. The distribution of specimens is summarized in Supplementary Table 1. Blood specimens were collected after aseptic preparation and cultured in trypticase soy broth (Becton Dickinson, MD, USA). Positive culture results were detected using the automated detection system (BD BACTEC™ FX; Becton Dickinson). Blood was drawn out from positive blood culture bottles onto blood plate (BP) agar for subculture (Becton Dickinson, MD, USA). Sputum specimens with adequate quality[1] were used. The respiratory specimens were inoculated on BP agar (Becton Dickinson), eosin methylene blue (EMB) agar (Becton Dickinson), CNA agar (Becton Dickinson), and chocolate agar (Becton Dickinson). Specimens obtained from the sterile cavity fluid were inoculated on BP, EMB, CNA, and chocolate agars, and into thioglycollate broth (Becton Dickinson). While positive growth was noted in thioglycollate broth, subculture was performed using BP agar. A semiquantitative culture method described by Maki et al. was used for testing the tip of implants.[2] Urine specimens were inoculated using a quantitative loop on BP and EMB agars. For specimens collected from wound, 1.2 mL of 0.9% saline was used for rinsing when the specimens were obtained using a swab. The rinsed saline was inoculated on BP, EMB, CNA, and chocolate agars; for pus collected from wound, the specimens were directly dropped on the agars and into thioglycollate broth. The agar and broth were incubated in a $CO_2$ incubator at 37°C for 18–24 hours. Single colonies grown on agar plates were picked for further analysis. *Enterococcus faecium* was identified based on colony morphology and matrix-assisted laser desorption ionization time-of-flight (MALDI-TOF) spectra (Bruker Daltonik GmbH, Bremen, Germany). The paper disc method was used to differentiate vancomycin-resistant *Enterococcus* from vancomycin-susceptible *Enterococcus* on the basis of Clinical and Laboratory Standards Institute guidelines M100.

**Supplementary Table 1.** Specimen Distribution

| **Total *E. faecium* Linkou (5717)** | **Blood (674)** | **Urinary tract (2818)** | **Sterile body fluid (1009)** | **Wound (1211)** | **Respiratory tract(3)** | **Others (2)** |
|---|---|---|---|---|---|---|
| S | 263(39.0%) | 1343(47.7%) | 664(65.8%) | 651(53.8%) | 1(33.3%) | 0(0%) |
| R | 411(61.0%) | 1475(52.3%) | 341(34.2%) | 560(46.2%) | 2(66.7%) | 2(100%) |
| **Total *E. faecium* Kaohsiung (2280)** | **Blood (205)** | **Urinary tract (988)** | **Sterile body fluid (338)** | **Wound (730)** | **Respiratory tract (1)** | **Others (18)** |
| S | 90(43.9%) | 524(53.0%) | 225(66.6%) | 219(30.0%) | 0(0%) | 0(0%) |
| R | 115(56.1%) | 464(47.0%) | 113(33.4%) | 511(70.0%) | 1(100%) | 18(100%) |

We only selected specimens of sufficient quantity, that is, blood, urinary tract, sterile body fluid, and wound, for building the vancomycin-resistant *E. faecium* (VRE*fm*) prediction model in this study.

**Analytical measurement by using MALDI-TOF mass spectrometry and data processing**

The analytical measurements of MALDI-TOF were conducted according to the manufacturer's instruction (Bruker Daltonik GmbH). Single colonies grown on agar were picked and smeared onto a MALDI steel target plate to form thin films. One microliter of 70% formic acid was applied on the films and dried at room temperature. One microliter matrix solution (50% acetonitrite containing 1% α-cyano-4-hydroxycinnamic acid and 2.5% trifluoroacetic acid) was then added on the films. The sample matrix was dried at room temperature before analysis using a mass spectrometer. MALDI-TOF was conducted using a Microflex LT mass spectrometer (Bruker Daltonik GmbH). Mass spectra were obtained under the following settings: linear positive mode; accelerating voltage +20 kV; and nitrogen laser frequency 60 Hz. Totally, 240 laser shots were hit on each sample spot for measurement. The Bruker Daltonics

Bacterial Test Standard was used for external calibration for the spectra. Flexanalysis 3.4 (Bruker Daltonik GmbH) was used for spectra processing. The Savitzky–Golay algorithm was set for spectra smoothing. The spectra baseline was subtracted using the top hat method. The signal-to-noise ratio threshold was set as 2. *E. faecium* was determined using Biotyper 3.1 (Bruker Daltonik GmbH) on the basis of processed spectra. All the spectra of the cases reached acceptable quality (log score ≥ 2, defined by manufacturer's instruction). Spectra ranging from 2000 to 20 000 Da were collected for further analysis.

All of the *E. faecium* isolates had a log score greater than 2 provided by Biotyper 3.1 (Bruker Daltonik GmbH, Bremen, Germany), which ensures the quality of MS spectra [3-5]. On this basis, we applied default preprocessing steps, including baseline subtraction, smoothing, and recalibration, to treat the MALDI-TOF spectra data (range from 2 000 to 20 000 Da) of each isolate by Flexanalysis 3.4 (Bruker Daltonik GmbH, Bremen, Germany) [6]. We then extracted the peaks with a high occurrence frequency as the predictor candidates through a binning-size method developed in a previous study,[4,7] which is illustrated in Supplementary Figure 2. The extracted peaks were then adjusted according to the alignment of *m/z* 4429, which was reported to be one of the conservative peaks for *E. faecium*[8,9] (Supplementary Figure 3). By aligning to the internal conservative peak, the possible shifting of spectra[4] observed could be adjusted.

**Binning method for extracting predictor candidates**

In a MALDI-TOF mass spectrometry (MS) spectra, the peaks were extracted and regarded as predictors for the construction of predictive models. In an initial scanning through all MS spectra, due to the isotope of various atoms, many peaks with a little bit difference of *m/z* values might be referred to as the same peptide species. The shifting problem of peaks is illustrated in Supplementary Figure 1. To deal with this problem among different spectra, the binning method was adopted to group large-scale peaks into a smaller number of "bins". Supplementary Figure 2 presents a schematic diagram of the binning method used in this study. The peaks located within the same bin are considered as the same feature. In the binning method, we evaluated various bin sizes (0-12 Da), and we adopted 10 Da as the bin size for the following experiments given the results of evaluation and the previous studies.[4,7] Moreover, on the basis of binning results, we aligned the peaks according to *m/z* 4429 to obtain more accurate *m/z* positions for the peaks. The peptide at *m/z* 4429 was reported to be fairly abundant in *E. faecium*.[8,9] Thus, we selected the peptide at *m/z* 4429 as the intrinsic internal control to adjust the other peaks (Supplementary Figure 3).

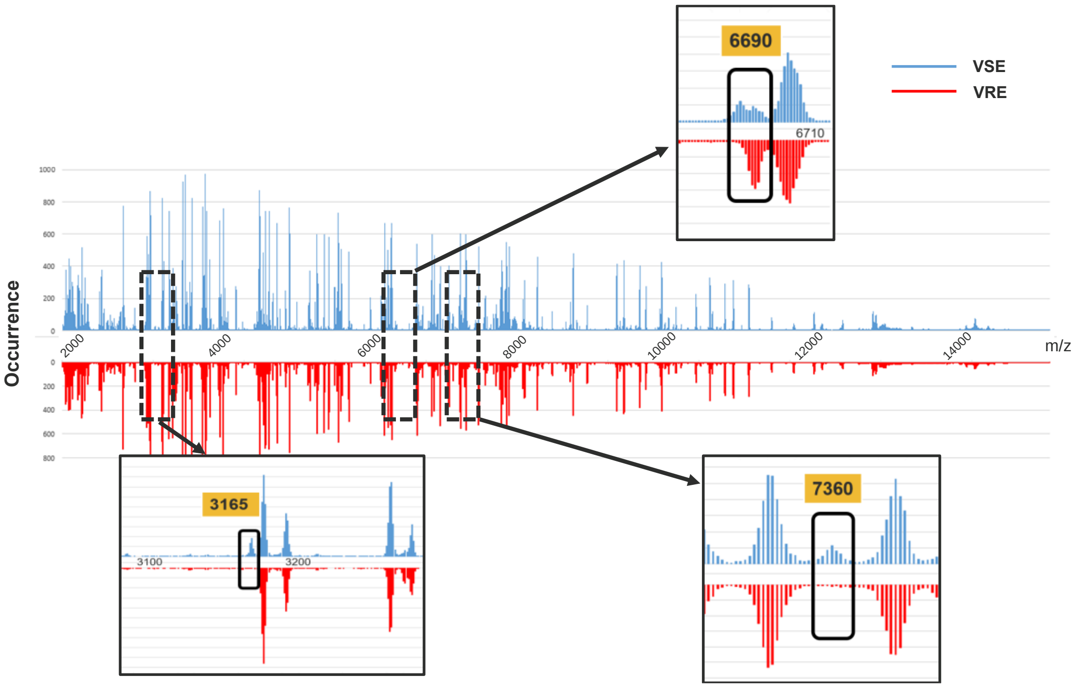

**Supplementary Figure 1.** An Example of MALDI-TOF MS Spectra to Illustrate the "Shifting Problem". Peptides of the same species do not appear at exactly the same *m/z* position. By contrast, the peptides of the same species would distribute in a normal distribution-like manner.

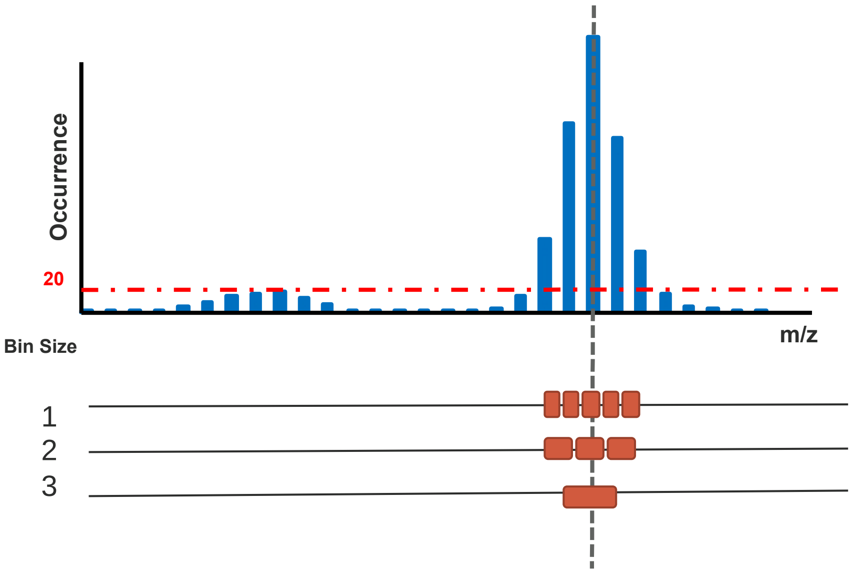


**Supplementary Figure 2.** Schematic Illustration of the Binning Method. In the original MS spectra, the peptides of the same species were located at several different *m/z* positions nearby. We used the binning method to classify these peaks into smaller groups. An example is illustrated: Occurrence frequency at a specific *m/z* larger than 20% of all cases is taken into calculation. When the bin size is 1, then 5 resulting features are obtained; when the bin size is 3, only 1 resulting feature is obtained.

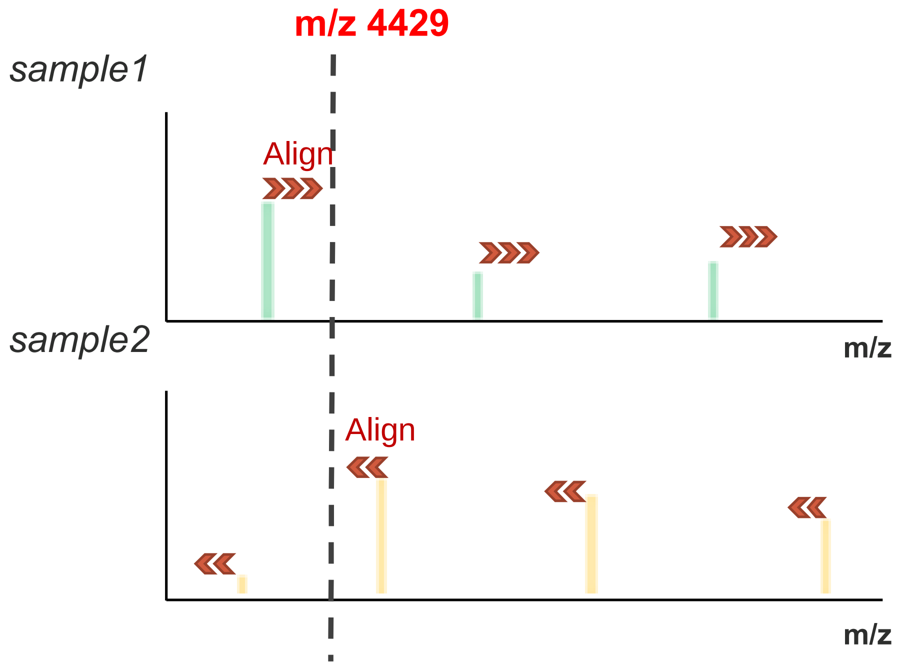


**Supplementary Figure 3.** Examples of Alignment to *m/z* 4429. We illustrated 2 examples of alignment: The upper one shows shifting to the right when the peak located closest to *m/z* 4229 is shifted from the left to right, and then, the other peaks on the spectra also shifted to the right. By contrast, the lower figure shows a case of shifting to the left when the peak closest to *m/z* 4429 shifted from right to left.

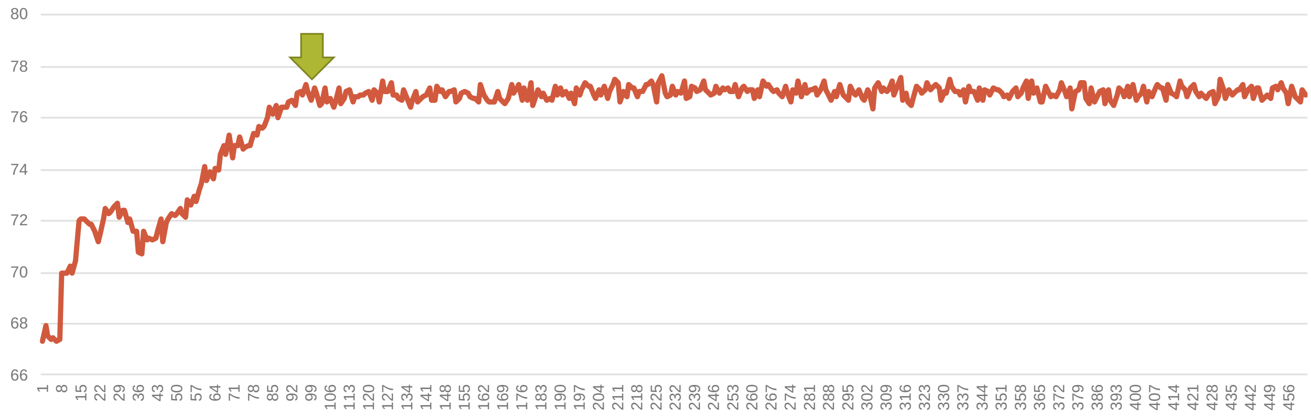

**Supplementary Figure 4.** Performance of ML Models Based on Different Number of Predictors. The performance reaches a plateau when the number of predictors is more than 100.

**Heat map**

On the basis of the chi-square scores of the predictor candidates (ie, peaks), we selected top 10 most critical predictive peaks (Supplementary Table 2) and plotted a heat map using hierarchical clustering. We took log of the intensity of the top 10 most critical predictive peaks, followed by z-score standardization. The hierarchical clustering was conducted based on Euclidean distance metric and average linkage. We produced the heat map by using pheatmap package on R software (version 3. 3. 3, R Foundation for Statistical Computing, http://www.r-project.org/).

**Random forest**

Random forest (RF), an ensemble learning method, is widely used for classification. More specifically, the ensemble learning approach combines multiple learning models to obtain an improved classification model and thus obtain better performance on prediction.[10] Additionally, the bootstrap aggregation (bagging) technique is considered for sampling the training data in RF. In other words, the bootstrap method samples the training tuples averagely with replacement, which means every selected tuple is likely to be re-added to the training set. In this study, we adopted the Weka toolkit[11] to construct RF classifiers based on various feature sets.

**K-Nearest Neighbors**

The nearest neighbor approach is an instance-based classifier used for determining the most similar instances, which were selected from all training data, to a given test instance, based on a distance function. Given a test instance, the most *k* similar instances are regarded as *k*-nearest neighbors (KNNs) of the test data, and the class assignment is determined in accordance with the proportion of KNNs.

Considering the training data and test data as the *n*-dimensional vectors in Gaussian space, the Euclidean distance function is usually applied to measure the distances between the test data and all training data. Given a test instance *t*, the Euclidean distance between *t* and a training instance *x* is defined as

$$\mathbf{d}(\boldsymbol{t},\boldsymbol{x}) = \sqrt{\sum\nolimits_{\boldsymbol{k}=\mathbf{1}}^{\boldsymbol{n}} (\boldsymbol{t}_{\boldsymbol{k}} - \boldsymbol{x}_{\boldsymbol{k}})^{\mathbf{2}}}$$

where *n* is the size of the feature set. After determining KNNs, the class labels of these k training instances might be inconsistent. A weighted distance voting method was used to conduct the class assignment for a test data. Class assignment C(*t*) of a test data *t* is determined by

$$\mathbf{C}(\boldsymbol{t}) = \underset{v}{\mathbf{argmax}} \sum_{x_i \in KNNs} \boldsymbol{w}_{\boldsymbol{i}} \times \boldsymbol{I}(\boldsymbol{x}_{\boldsymbol{i}}) \qquad \begin{cases} \boldsymbol{I}(\boldsymbol{x}_{\boldsymbol{i}}) = \mathbf{1} \;\; \boldsymbol{if} \;\; \boldsymbol{v} = \boldsymbol{class} \;\; \boldsymbol{label} \;\; \boldsymbol{of} \;\; \boldsymbol{x}_{\boldsymbol{i}} \\ \boldsymbol{I}(\boldsymbol{x}_{\boldsymbol{i}}) = \mathbf{0} \;\; \boldsymbol{if} \;\; \boldsymbol{v} \neq \boldsymbol{class} \;\; \boldsymbol{label} \;\; \boldsymbol{of} \;\; \boldsymbol{x}_{\boldsymbol{i}} \end{cases}$$

where *v* is the class label and $\boldsymbol{w}_{\boldsymbol{i}}$ is the weighted value of the class label of $\boldsymbol{x}_{\boldsymbol{i}}$ in KNNs. For a binary classification between VRE*fm* and vancomycin-susceptible *E. faecium* (VSE*fm*) samples, the positive and negative training instances were represented as *n*-dimensional vectors with class labels +1 and −1, respectively. The testing data without class labels are classified into +1 or −1 based on the *k* nearest training samples. In the KNN classifier, various values of *k* were examined to find the best cutoff with best performance.

**Support Vector Machine**

This study involved a binary classification of VRE*fm* and VSE*fm* spectra. The positive (VRE*fm*) and negative (VSE*fm*) spectra were labeled as +1 and −1, respectively, for the 2 classes. The training dataset is $X = \{x^t, c^t\}$ where $c^t = +1$ if $x^t \in$ positive dataset and $c^t = -1$ if $x^t \in$ negative dataset. This study attempted to identify *w* and $w_0$ such that

$w^T x^t + w_0 \geq +1 \quad for \;\; c^t = +1$ and $w^T x^t + w_0 \leq -1 \quad for \;\; c^t = -1$,

which can be rewritten as

$$c^t(w^T x^t + w_0) \geq +1$$

This formula could be used to estimate the optimal separating hyperplane that can maximize the margin between 2 classes.[12] The distance of $x^t$ to the discriminating hyperplane is

$$\frac{|w^T x^t + w_0|}{||w||}$$

and we would like the distance to be higher than a specific value *h*:

$$\frac{c^t(w^T x^t + w_0)}{||w||} \geq h, \forall t \ \ and \ \ c^t \in \{+1, -1\}$$

The support vector machine (SVM) is an advanced algorithm used to identify a hyperplane between 2 classes with a maximum margin based on an *n*-dimensional vector space.[12] In an attempt to maximize *h*, however, an unlimited number of possible values could be elucidated by tuning *w*. Hence, $h||w||$ was defined as one and $||w||$ was minimized using the following equation[13]:

$$min \frac{1}{2}||w||^2 \ \ subject \ \ to \ \ c^t(w^T x^t + w_0) \geq +1, \forall t$$

In this work, SVM could be adopted to determine a hyperplane for discriminating between VRE*fm* and VSE*fm* samples with a maximal margin in a vector space containing *n* dimensions (size of the feature set). The mass-to-charge ratio values of spectra were represented as a numeric vector in an n-dimensional vector space, which are the input values for SVM. A famous SVM public resource, called LIBSVM,[14] was downloaded and installed in our computing server for an iterative training of multiple SVMs in accordance with various feature sets. With ML, if the best discriminant is nonlinear, instead of enabling a nonlinear model, we could map all *n*-dimensional vectors to a new vector space with higher dimension *m*, where $m > n$, based on nonlinear kernel functions. As demonstrated in previous methods,[15-19] the radial basis function (RBF) has been typically chosen as the specified kernel function in SVM models. The RBF function was given as follows:

$$K(x^t, x) = \exp\left[-\frac{||x^t - x||^2}{2s^2}\right]$$

where $x^t$ is the center and *s* is the radius, which should be provided by the programmer. With

LIBSVM, cost (*c*) and gamma (*r*) are 2 supporting parameters used to optimize the radius of the kernel function and softness of the hyperplane, respectively. To achieve the feasible values of gamma (*r*) and cost (*c*) in model learning, an optimization program, written in Python, was provided by LIBSVM.

**Supplementary Table 2.** Top 100 Important Predictors Calculated Using Chi-Square

| Predictor (m/z) | X-squared | P-value |
|---|---|---|
| **6690** | 749.6721 | 4.73E-165 |
| **6603** | 680.4819 | 5.25E-150 |
| **3302** | 520.7866 | 2.85E-115 |
| **3165** | 461.9951 | 1.77E-102 |
| **6342** | 451.5601 | 3.30E-100 |
| **3645** | 450.855 | 4.70E-100 |
| **3172** | 412.4703 | 1.06E-91 |
| **6356** | 391.5542 | 3.80E-87 |
| **7360** | 375.9846 | 9.32E-84 |
| **3681** | 344.9242 | 5.40E-77 |
| **7289** | 317.5502 | 4.95E-71 |
| **3655** | 299.2888 | 4.71E-67 |
| **6661** | 271.7419 | 4.73E-61 |
| **6528** | 254.4583 | 2.77E-57 |
| **6512** | 246.2366 | 1.72E-55 |
| **12711** | 233.6196 | 9.68E-53 |
| **3740** | 211.9791 | 5.08E-48 |

| | | |
|---|---|---|
| **7306** | 208.538 | 2.86E-47 |
| **6741** | 199.5891 | 2.57E-45 |
| **3651** | 197.2695 | 8.24E-45 |
| **3900** | 175.462 | 4.75E-40 |
| **6361** | 174.2589 | 8.69E-40 |
| **10625** | 167.6077 | 2.46E-38 |
| **3870** | 164.2027 | 1.37E-37 |
| **5949** | 145.6891 | 1.52E-33 |
| **6328** | 137.147 | 1.12E-31 |
| **13482** | 133.0305 | 8.90E-31 |
| **3264** | 130.0682 | 3.96E-30 |
| **6631** | 126.649 | 2.22E-29 |
| **7385** | 122.6643 | 1.65E-28 |
| **7310** | 120.8941 | 4.03E-28 |
| **3884** | 110.0463 | 9.57E-26 |
| **3306** | 99.5212 | 1.94E-23 |
| **6748** | 94.5692 | 2.37E-22 |
| **10629** | 90.9076 | 1.51E-21 |
| **2430** | 86.3685 | 1.49E-20 |

| | | |
|---|---|---|
| **12715** | 84.3901 | 4.06E-20 |
| **5935** | 77.9294 | 1.07E-18 |
| **6665** | 76.8498 | 1.84E-18 |
| **7413** | 68.7908 | 1.09E-16 |
| **3906** | 68.5522 | 1.24E-16 |
| **2720** | 68.3585 | 1.36E-16 |
| **3316** | 60.3783 | 7.83E-15 |
| **6308** | 59.4367 | 1.26E-14 |
| **3256** | 59.0654 | 1.53E-14 |
| **6333** | 58.9954 | 1.58E-14 |
| **5478** | 52.5241 | 4.25E-13 |
| **12720** | 50.6321 | 1.11E-12 |
| **3875** | 50.3945 | 1.26E-12 |
| **4857** | 48.9752 | 2.59E-12 |
| **7743** | 48.5973 | 3.14E-12 |
| **12724** | 48.5585 | 3.21E-12 |
| **7364** | 48.0801 | 4.09E-12 |
| **6607** | 45.4681 | 1.55E-11 |
| **6635** | 41.967 | 9.28E-11 |

| | | |
|---|---|---|
| **7753** | 41.6587 | 1.09E-10 |
| **5768** | 40.7215 | 1.76E-10 |
| **3518** | 40.3004 | 2.18E-10 |
| **5313** | 39.0606 | 4.11E-10 |
| **6493** | 38.438 | 5.65E-10 |
| **4831** | 37.1236 | 1.11E-09 |
| **2967** | 37.0867 | 1.13E-09 |
| **6460** | 36.16 | 1.82E-09 |
| **9714** | 34.0623 | 5.34E-09 |
| **4573** | 33.8151 | 6.06E-09 |
| **3694** | 32.8717 | 9.84E-09 |
| **6477** | 32.6426 | 1.11E-08 |
| **7034** | 32.3444 | 1.29E-08 |
| **7418** | 31.5142 | 1.98E-08 |
| **4469** | 31.0475 | 2.52E-08 |
| **3013** | 30.8891 | 2.73E-08 |
| **3915** | 30.3285 | 3.65E-08 |
| **6752** | 30.3066 | 3.69E-08 |
| **9773** | 30.2625 | 3.77E-08 |

| | | |
|---|---|---|
| **10957** | 29.5373 | 5.49E-08 |
| **9953** | 29.1471 | 6.71E-08 |
| **13385** | 29.1427 | 6.72E-08 |
| **7330** | 29.0184 | 7.17E-08 |
| **10941** | 28.9438 | 7.45E-08 |
| **7445** | 28.6333 | 8.75E-08 |
| **3337** | 28.5984 | 8.91E-08 |
| **3321** | 27.3422 | 1.70E-07 |
| **5038** | 25.7023 | 3.98E-07 |
| **3724** | 25.4256 | 4.60E-07 |
| **5220** | 25.0366 | 5.63E-07 |
| **12728** | 24.7339 | 6.58E-07 |
| **4861** | 24.3722 | 7.94E-07 |
| **7275** | 24.3091 | 8.20E-07 |
| **5387** | 23.9425 | 9.93E-07 |
| **10952** | 23.9008 | 1.01E-06 |
| **5485** | 23.6539 | 1.15E-06 |
| **6346** | 23.6241 | 1.17E-06 |
| **4844** | 23.5285 | 1.23E-06 |

| | | |
|---|---|---|
| **6105** | 23.485 | 1.26E-06 |
| **5383** | 23.3773 | 1.33E-06 |
| **10075** | 22.8121 | 1.79E-06 |
| **8031** | 22.4537 | 2.15E-06 |
| **2449** | 22.0113 | 2.71E-06 |
| **6818** | 21.9906 | 2.74E-06 |
| **3027** | 21.2967 | 3.93E-06 |

**Supplementary Table 3(a).** Comparison of AUROCs Between Different Algorithms With Different Validation Methods

| | 5-fold CV | Time-wise internal validation | External validation |
|---|---|---|---|
| KNN-RF | $1.37\times10^{-45}$ | $4.94\times10^{-11}$ | $1.24\times10^{-15}$ |
| KNN-SVM | $7.16\times10^{-24}$ | $6.49\times10^{-7}$ | $2.49\times10^{-7}$ |
| RF-SVM | $3.33\times10^{-8}$ | 0.0310 | 0.0001 |

**Table 3(b).** Comparison of Accuracies Between Different Algorithms With Different Validation Methods, and *P* Values of Cochran's Q Test

| Datasets | RF | SVM | KNN | p-value |
|---|---|---|---|---|
| 5-fold CV | 0.7769 (0.7660, 0.7878) | 0.7610 (0.7499, 0.7721) | 0.7248 (0.7131, 0.7364) | $2.20\times10^{-16}$ |
| Time-wise internal validation | 0.7840 (0.7640, 0.8039) | 0.7815 (0.7615, 0.8016) | 0.7228 (0.7011, 0.7445) | $1.72\times10^{-12}$ |
| External validation | 0.7855 (0.7687, 0.8024) | 0.7781 (0.7610, 0.7951) | 0.7355 (0.7174, 0.7536) | $5.15\times10^{-11}$ |

**Table 3(c).** Comparison of Sensitivities Between Different Algorithms With Different Validation Methods, and *P* Values of Cochran's Q Test

| Datasets | RF | SVM | KNN | p-value |
|---|---|---|---|---|
| 5-fold CV | 0.8054 (0.7951, 0.8517) | 0.7826 (0.7719, 0.7934) | 0.7873 (0.7767, 0.7980) | 0.0038 |
| Time-wise internal validation | 0.8153 (0.7965, 0.8341) | 0.8415 (0.8238, 0.8592) | 0.7491 (0.7281, 0.7702) | $1.50\times10^{-12}$ |
| External validation | 0.7791 (0.7620, 0.7961) | 0.7954 (0.7789, 0.8120) | 0.8044 (0.7881, 0.8207) | 0.0265 |

**Table 3(d).** Comparison of Specificities Between Different Algorithms With Different Validation Methods, and *P* Values of Cochran's Q Test

| Datasets | RF | SVM | KNN | p-value |
|---|---|---|---|---|
| 5-fold CV | 0.7497 (0.7384, 0.7609) | 0.7403 (0.7289, 0.7517) | 0.6649 (0.6526, 0.6772) | $2.20\times10^{-16}$ |
| Time-wise internal validation | 0.7477 (0.7266, 0.7688) | 0.7120 (0.6900, 0.7340) | 0.6922 (0.6698, 0.7146) | 0.0002 |
| External validation | 0.7930 (0.7764, 0.8096) | 0.7580 (0.7405, 0.7756) | 0.6560 (0.6365, 0.6755) | $2.20\times10^{-16}$ |

**Table 3(e).** Pairwise McNemar's Test for Sensitivity, Specificity, and Accuracy Between Different Algorithms With Different Validation Methods

| Comparison | Adjusted p-value of comparing two sensitivities | Adjusted p-value of comparing two specificities | Adjusted p-value of comparing two accuracies |
|---|---|---|---|
| (a) 5-fold CV | | | |
| KNN-RF | 0.0284 | $1.08\times10^{-26}$ | $7.86\times10^{-21}$ |

| | | | |
|---|---|---|---|
| KNN-SVM | 0.5640 | $5.58\times10^{-20}$ | $5.62\times10^{-10}$ |
| RF-SVM | $8.70\times10^{-5}$ | 0.0909 | $4.14\times10^{-5}$ |
| (b) Time-wise internal validation | | | |
| KNN-RF | $4.44\times10^{-6}$ | $4.28\times10^{-4}$ | $1.14\times10^{-8}$ |
| KNN-SVM | $1.10\times10^{-9}$ | 0.2100 | $7.46\times10^{-8}$ |
| RF-SVM | $3.76\times10^{-3}$ | $3.45\times10^{-4}$ | 0.7050 |
| (c) External validation | | | |
| KNN-RF | 0.0366 | $1.25\times10^{-21}$ | $1.36\times10^{-8}$ |
| KNN-SVM | 0.3830 | $3.00\times10^{-13}$ | $8.20\times10^{-7}$ |
| RF-SVM | 0.0618 | $3.94\times10^{-4}$ | 0.2350 |
